# Acute exposure to groundwater contaminants mixture of nitrate, atrazine and imidacloprid impacts growth kinetics of poultry cecal microbiomes and significantly decreases Caco-2 cell viability

**DOI:** 10.1101/2025.06.15.659797

**Authors:** Chamia C. Chatman, Erica L.-W. Majumder

## Abstract

Atmospheric deposition, and agricultural runoff or erosion events have substantially contributed to groundwater pollution throughout the USA. This can become troublesome in states like Wisconsin where 68% of the population rely on groundwater for their drinking water source. As such, exposome research must account for the complexity and frequency of environmental exposures. This study aimed to elucidate chemical-biological interactions and adverse outcome pathways associated with an environmentally relevant mixture of agricultural chemicals detected in Wisconsin groundwater via *in vitro* and *in silico* methodologies. Using *in vitro* models, we determined that a ternary mixture of environmentally relevant concentrations of nitrate, atrazine and imidacloprid resulted in an overt decline in growth rate to the poultry cecal microbiome compared to each chemical singularly. Further, there was a decrease in Caco-2 cell viability in various two-chemical combinations. *In silico* methods analyzed contaminants detected in Wisconsin groundwater wells from across the state and prioritized two groundwater wells as potential for health concerns. Prioritized chemicals in these groundwater wells were linked to nine gene targets and several adverse outcome pathways. In all, the results demonstrated that there is chemical-biological interaction between these model organisms agricultural and chemical mixtures at real world exposure concentrations.

**Highlights:**

- *in silico* methods were able to predict potential adverse effects for communities utilizing these groundwater wells
- 8 out of 9 chemicals prioritized with *in silico* tools were herbicides
- A ternary mixture of nitrate, imidacloprid and atrazine resulted in a decline in growth rate for poultry cecal microbiome
- Caco-2 cells significantly impacted by two-chemical combinations but not ternary mixtures

## 1. Introduction

Within the United States of America (USA), nearly half of the land is utilized for agriculture [1]. To date, agricultural operations, a type of nonpoint source pollution, have substantially contributed to the contamination of groundwater [2,3]. Point source pollution is defined as pollutants from a fixed or confined location like wells, pipes, containers, or vessels [4]. Annually in the USA approximately a half million tons of pesticides, 12 million tons of nitrogen, and 4 million tons of phosphorus fertilizer are applied annually to crops [1]. Consequently, atmospheric deposition, and agricultural runoff or erosion events have substantially contributed to groundwater pollution throughout the USA [3,5]. This can become a public health concern in states such as Wisconsin where approximately 68% of the population relies on groundwater as a primary drinking water source [6]. Likewise, livestock production systems in Wisconsin utilize mostly groundwater [7].

In Wisconsin, agriculture is a major economic sector which contributes over $100 billion annually to the state’s economy [8]. A common practice has been to apply pesticides, herbicides and fertilizers annually. Nitrogen, a primary macronutrient, was applied to an estimated 88% of corn crops planted in Wisconsin during the 2021 crop year [9]. The application of nitrate fertilizers has directly contributed to nearly 90% of total nitrate contamination in Wisconsin groundwater [6]. In more rural areas of the state, more than 20% of the groundwater wells contain nitrate concentrations above the groundwater standard of 10 mg/L [6]. Consumption of food or water contaminated with nitrate has been associated with cancer, fetal malformations, and spontaneous abortions [6,10]. Methemoglobin concentrations in infants have been strongly correlated to nitrate consumption [10]. Methemoglobinemia occurs when nitrate is reduced to nitrite via bacteria in saliva or the stomach which then attaches to hemoglobin and forms methemoglobin [10]. Unlike hemoglobin, methemoglobin is not capable of binding to oxygen and therefore cannot carry oxygen through the body [10,11]. As a result, the US EPA set a standard of 10 mg/L to protect more vulnerable populations like children and pregnant women [10].

During this same crop growing year, atrazine was the predominate herbicide used. Specifically, 95% of corn acres planted in Wisconsin had an herbicide active ingredient applied, of which atrazine contributed 64% [9]. Atrazine is a persistent chlorotriazine herbicide which inhibits the photosynthetic pathway of several broadleaf weeds [12]. Exposure to atrazine has been demonstrated to alter reproductive development [13–15], disrupt the endocrine system [16,17], and induce inflammation in the brain [18,19]. Considering the biological effects noted in *in vivo* and *in vitro* atrazine exposure studies, it is concerning that this chemical has been detected in human urine samples [20,21]. In 1991, the Atrazine Rule was developed as a preventative measure to reduce atrazine contamination in groundwater [22]. Prohibited atrazine use now cover 25 counties or 1.2 million acres [23]. Monitoring of groundwater has demonstrated that atrazine concentrations have decreased in prohibited use areas but continues to be groundwater contaminant of concern [23].

Alongside nitrate and atrazine, imidacloprid was the other dominant chemical applied to corn crops in Wisconsin in the 2021 crop year. Imidacloprid was detected in 34% (31 of 951 samples) of samples analyzed from monitoring wells near agricultural fields in Wisconsin [24]. Imidacloprid is a neonicotinoid insecticide which acts on the acetyl neuronal acetylcholine receptors in insects thus causing central nervous system disorders [12]. Application of the agricultural chemical has been documented to have off-target effects in various species, most notably honeybees [25,26]. This resulted in the ban of imidacloprid and other neonicotinoids in the European Union in 2018 [27].

There are several knowledge gaps regarding chemical mixtures detected in groundwater. Mainly due to the detection of chemicals without groundwater standards or health advisory levels. This was observed in the 2023 Wisconsin Groundwater Quality report where 13 out of 29 pesticides had no groundwater standards [28]. The lack of groundwater standards further exacerbates public concerns regarding groundwater contamination as there are unknown health risks following exposure. In addition, chemicals with known groundwater standards tend to be evaluated on an individual basis and not as a part of an ecologically relevant mixture [29]. There have been improvements to chemical mixture risk assessments. However, a key limitation is the lack of directly measured mixtures and inconsistent toxicological data for individual chemicals [30], both of which are essential for developing risk assessment frameworks. To provide more accurate chemical risk assessments, environmental concentrations of chemical mixtures need to be evaluated. This will provide more realistic data as to if and how chemical mixtures led to adverse health impacts to hosts.

Here, we aimed to elucidate chemical-biological interactions and adverse outcome pathways associated with an environmentally relevant mixture of agricultural chemicals detected in Wisconsin groundwater. Nitrate, atrazine and imidacloprid were the selected chemicals of interest based on their frequency of cooccurrence in groundwater wells and chemical concentrations for each exceeding either an enforcement standard or preventative action limit in the 2021 Department of Agriculture, Trade, and Consumer Protections (DATCP) Targeted Sampling Report [31]. We hypothesized that exposure to nitrate, atrazine, and imidacloprid would result in a more toxic response compared to exposure to each chemical individually. We further hypothesized that impact to this chemical mixture would vary between the gut microbiome and gastrointestinal epithelial cells. To test these hypotheses and elucidate the impact oral ingestion of contaminated groundwater may have on human and livestock health, we exposed gastrointestinal tract microbiome and epithelial cells to nitrate, atrazine and imidacloprid singularly, in a two-chemical combination or as ternary mixtures. We also incorporated *in silico* tools to determine if this chemical concentration data correlated to chemical-biological effects. In this study, we demonstrate that it is imperative that toxicology research evaluates chemical mixtures at environmentally relevant concentrations as observed concentrations were well below the cytotoxicity levels but still disturbed the growth and physiology of the cells that could contribute to negative health outcomes as a chronic exposure.

## 2. MATERIALS AND METHODS

### 2.1 Chemical prioritization using *toxEval* and *ToxMixtures*

Data from the 2021 DATCP Targeted Sampling Report was utilized for chemical prioritization [31]. This report summarized water quality analyses conducted on groundwater well samples collected from private groundwater wells in agricultural areas throughout Wisconsin (16 counties, N = 104 samples) (Figure 1). The sampling period was conducted in June and September 2021. Additional information regarding the analytical testing is provided in the 2021 DATCP Targeted Sampling Report [31]. The chemical abstract numbers (CAS), and chemical concentration data were compiled for each chemical detected and added to the metadata file (Supplemental Tables 1-3). Chemical mixtures of interest and their associated gene targets and adverse outcome pathways were then determined using the R-packages *toxEval* [32] *and* ToxMixtures [33].

**Figure 1.**
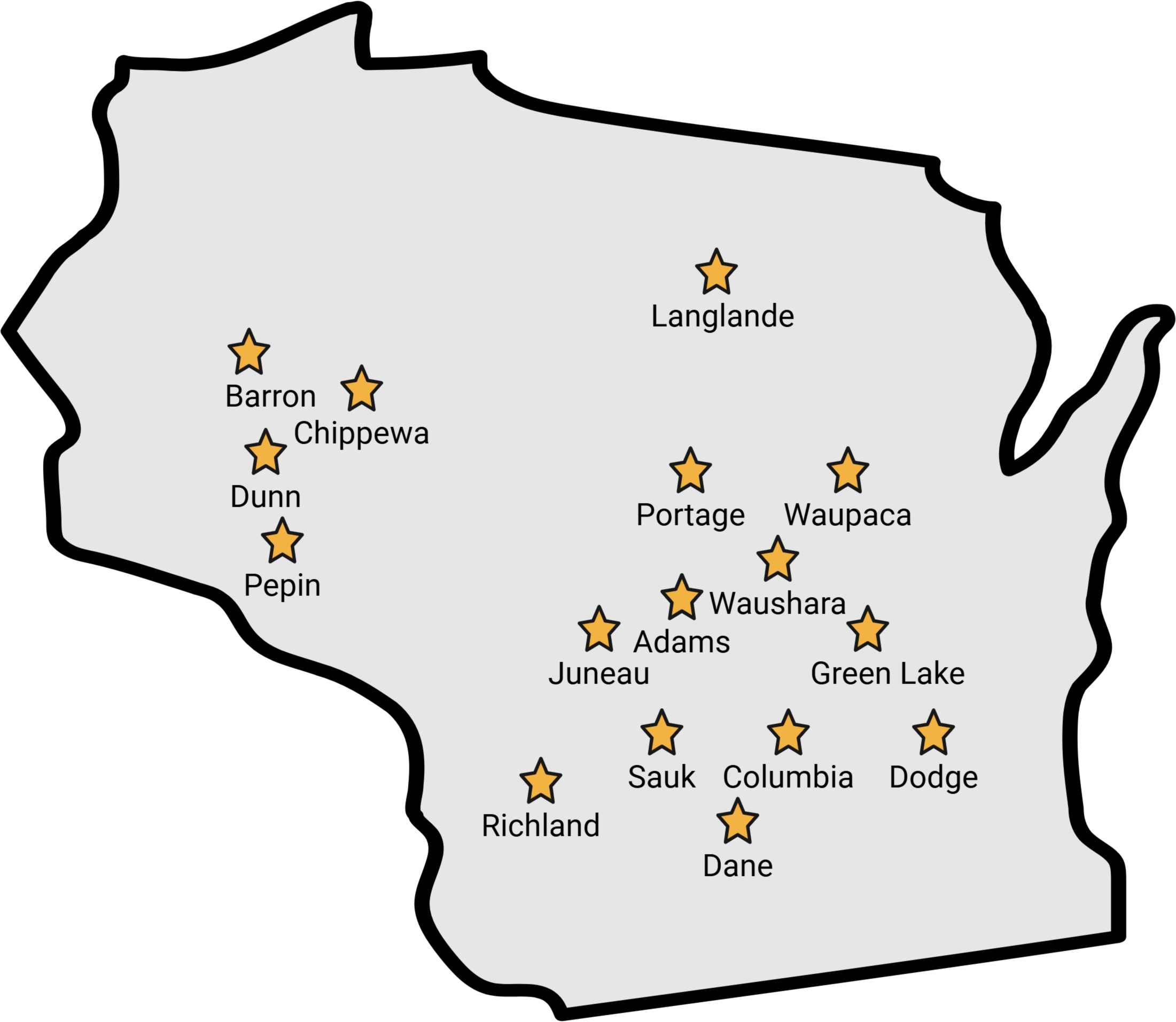
The 2021 DATCP Targeted Sampling Report evaluated groundwater samples from 16 counties in Wisconsin (gold stars) (N = 104).

We calculated the exposure-activity ratios (EAR) for detected chemicals using the R-package *toxEval* [32] to prioritize agricultural chemicals of potential biological concern. EARs are defined as the ratio of the measured chemical concentration and the activity concentration at cutoff (ACC) [34]. The ACC is used in the EAR calculation instead of AC50 because the ACC is less prone to violating assumptions about the relative potency of a given chemical [34]. An EAR threshold of 10^-3^ is used as this value has been demonstrated to be a level of potential concern based on comparison with established water quality benchmarks [35]. An EAR threshold of 10^-3^ has also been shown to prioritize an analogous list of chemicals in water quality data as a toxicity quotient (TQ) = 0.1 [36]. TQ is the ratio of the measured concentration of a chemical (µg/L) and the benchmark concentration (µg/L) [36]. An EAR ≥ 1 indicates that the measured concentration is greater than the ToxCast endpoint concentration [37]. This is possible to derive because *toxEval* and ToxMixtures calculate the EAR and correspond the water quality data to ToxCast assay endpoints (Version 3.5) [32,35]. ToxCast (Toxicity Forecasting) is a program developed by the United States Environmental Protection Agency (US EPA) which makes high-throughput *in vitro* data publicly available for prioritizing and characterizing chemicals [38]. Thus, potential chemical-biological effects can be determined by comparing the water quality data to the ToxCast assays.

ToxMixtures was subsequently used to calculate EAR_mixture_, EAR_gene_ and EAR_AOP_. The EAR_mixture_ is defined as the ∑EAR for chemicals detected within the same ToxCast assay endpoint [36]. The EAR_gene_ is the ∑EAR for all chemicals detected within the same ToxCast assay endpoint that related to common gene targets [33]. The EAR_AOP_ is defined as the ∑EAR of chemicals active on ToxCast endpoints (i.e., each assay-chemical combination) [33,36].

### 2.2 Chemicals and Reagents

The highest reported concentrations of each chemical from the 2021 DATCP Targeted Sampling Report were used for cell proliferation and viability experiments. Chemicals included atrazine (ASTATACH, Lot: P151-04287), imidacloprid (Thermo Scientific, Lot: A0450203), and nitrate (AccuStandard, Lot: 222105041). Atrazine was first diluted in DMSO then final dilutions were done in Dulco’s modified Eagle’s medium or 1x Luria-Broth (Difco; Lot:244620) depending on the experiment conducted. Nitrate and Imidacloprid were first diluted in autoclaved Milli-Q water then final dilutions were done in Dulbecco’s modified Eagle’s medium (ThermoFisher, ref: 11885-084) or 1x Luria-Broth (Difco; Lot:244620) depending on the experiment conducted.

### 2.3 Agricultural chemical mixtures

The DATCP mixture represents chemical concentrations like those detected in the 2021 DATCP Targeted Sampling Report. The equipotent and equivalent mixtures were selected based on preliminary experiments that used the methods described in section 2.5. Preliminary ternary exposure experiments indicated that equivalent concentrations of 100 ppb and 10 ppb elicit the greatest inhibition to poultry cecal microbiomes samples (Figure S1). To elicit a stronger phenotype, 100,000 ppb (a 1,000-fold increase) was used for each chemical in the equivalent mixture. The results of preliminary checkerboard experiments highlighted that lower doses of nitrate combined with higher concentrations of atrazine would lead to more growth inhibition in poultry cecal microbiomes (Figure S2). To further examine this, 10,000 ppb atrazine and 10,000 ppb imidacloprid (100-fold increase based on ternary exposure) and 1,000 ppb nitrate (10-fold increase based on ternary exposure) were used for the equipotent dose.

All two-chemical screening experiments were conducted with 3,000 ppb to 0 ppb (10-fold dilution) of nitrate, atrazine and imidacloprid. Two-chemical combinations included either imidacloprid and nitrate, imidacloprid and atrazine or nitrate and atrazine. The starting concentration of 3,000 ppb is a 30-fold increase of the concentrations used in preliminary ternary exposure studies (Figures S1) The ternary mixtures were as follows: 1) the environmental mixture (35,000 ppb nitrate + 1.7 ppb atrazine + 0.58 ppb imidacloprid), 2) the equipotent mixture (1,000 ppb nitrate + 10,000 ppb atrazine + 10,000 ppb imidacloprid), and 3) the equivalent mixture (100,000 ppb nitrate + 100,000 ppb atrazine + 100,000 ppb imidacloprid) (Table 1). Each chemical used for the DATCP mixture was also evaluated individually.

**Table 1.** Composition of agricultural chemical mixtures used in this study.

|  |  |  |  |
| --- | --- | --- | --- |
|  | **Nitrate**<br>**Concentration**<br>**(ppb)** | **Atrazine**<br>**Concentration (ppb)** | **Imidacloprid**<br>**Concentration**<br>**(ppb)** |
| DATCP<br>Mixture | 35,000 | 1.7 | 0.58 |
| Equipotent<br>Mixture | 1,000 | 10,000 | 10,000 |
| Equivalent<br>Mixture | 100,000 | 100,000 | 100,000 |

### 2.4 Caco-2 cell culture

The human colon adenocarcinoma cell lines (passage 6) were obtained from the laboratory of Dr. Ophelia Venturelli at the University of Wisconsin-Madison. Caco-2 cells were cultured in Dulbecco’s modified Eagle’s medium (ThermoFisher, ref: 11885-084), supplemented with 10% heat-inactivated fetal bovine serum (Gibco, Lot: 2687344RP), and 1% penicillin-streptomycin solution (Gibco, Lot: 2585652) in a humidified atmosphere (5% CO_2_, 95% air, 37°C). The cells were grown in T75 flasks under standard conditions until 70-80% confluent. Cells were harvested by trypsin-EDTA treatment. Cells were seeded at 1.6 x 10^4^ cells/mL in 96-well Falcon tissue culture plates (Ref: 353072). Cells from passages 8 through 9 were used for the experiments. The formation of polarized Caco-2 cell monolayers was not confirmed. Medium was changed every 2-3 days. Cells were used 24 h after plating into the 96-wll plates. Confluent cells were exposed to either one chemical, ternary mixtures or a two-chemical combination.

### 2.5 Broiler cecal culture preparation

Cecal digesta was collected from a previous published study and diluted following methods by Chatman et al (2024) [39]. In brief, ceca from untreated broilers were removed at the ileal-cecal junction and placed in sterile collection bags. Subsequent methods were performed under anaerobic conditions in a Coy (Coy Laboratory Products, Grass Lake, Michigan, USA) anaerobic chamber with atmosphere containing 5% O_2_, 10% CO_2_, and 85% N_2_). Then 0.1 g of cecal content from the proximal end of each cecum was collected into sterile 2 mL centrifuge tubes and resuspended in 900 µL of Anaerobic Dilution Solution (0.45 g/L K_2_HPO_4_, 0.45 g/L KH_2_PO_4_, 0.45 g/L (NH_4_)_2_SO_4_, 0.9 g/L NaCl, 0.1875 g/L MgSO_4_-7H_2_O, 0.12 g/L CaCl_2_-2H_2_O, 0.0001% resazurin, 0.05% cysteine-HCl, and 0.2% NaCO_3_). Resuspended cecal samples were subsequently diluted 1:3,000 in ADS. Then 0.5 mL of a 50% glycerol stock solution was added to each 1 mL aliquot of diluted cecal digesta. Glycerol stocks of broiler cecal cultures were stored at -80°C until liquid cultures were started.

The day prior to the experiment, a 1.5 mL broiler cecal glycerol aliquot was added to 10 mL of 1x Luria-Broth (Difco; Lot:244620) under anaerobic conditions. The culture was capped with a rubber stopper and aluminum crimp to maintain anaerobic conditions in the glass tube. The culture was then incubated overnight at 42°C with continuous shaking. The following day absorbance was measured at 595 nm to ensure bacterial growth. Bacterial growth was confirmed if the culture was turbid and the OD_600nm_ was approximately 0.1.

### 2.5 Evaluation of broiler cecal growth kinetics over 24-hour exposure

A preliminary growth curve experiment using glycerol stocks of poultry cecal samples stored from previous published experiments was conducted to determine ternary mixture concentrations. The growth kinetic experiment was performed with a checkerboard method to assess A) a two-chemical combination of atrazine and imidacloprid (300 to 0.16 ppb), B) a two-chemical combination of nitrate and atrazine (300 to 0.16 ppb) and C) ternary mixtures of nitrate, atrazine and imidacloprid at 100 µg/mL to 0.1 µg/mL (10-fold serial dilution). Chemical dilutions were done with autoclave water. A 150 µL aliquot of the cecal culture was added to each well in a 96-well plate followed by the addition of 50 µL of either ternary mixture (1:1:1 v/v) and subsequently mixed by pipetting up and down. In the anaerobic chamber, the 96-well plate was inserted into the Stratus microplate reader (Cerrillo, Charlottesville, Virginia, USA). The microplate reader was then immediately removed from the anaerobic chamber and placed in an anaerobic jar (Mitsubishi™ AnaeroPack-Anaero Gas Generator, ThermoScientific™, Waltham, MA, USA) and incubated at 37°C for 24 h. The absorbance (OD_600 nm_) was recorded every 30 min for 24 h.

Growth kinetics was also assessed for the ternary mixtures and each chemical at the reported concentrations from the 2021 DATCP Targeted Sampling Report. In the anaerobic chamber, 100 µL of the cecal culture was added to each well in a 96-well plate followed by the addition of 100 µL of either ternary mixture (1:1:1 v/v) and subsequently mixed by pipetting up and down. In the anaerobic chamber, the 96-well plate was inserted into the Stratus microplate reader (Cerrillo, Charlottesville, Virginia, USA). The microplate reader was then immediately removed from the anaerobic chamber and placed into an Anoxomat Anaerobic Jar System (Advanced Instruments, Norwood, Massachusetts, USA). The Anoxomat was calibrated to an anaerobic environment. Once the system indicated the jar was anaerobic, the Anoxomat was incubated at 42°C with continuous shaking for 24 h. The absorbance (OD_600 nm_) was recorded every 30 min for 24 h. This experiment was performed 3 times with 4 replicates per condition. The average log absorbance values from 3 independent experiments were plotted in GraphPad Prism (Version 10.3.1).

### 2.6 Caco-2 cell viability using the MTT assay

Caco-2 cells, a human-derived colon adenocarcinoma cell line, have historically been used for pesticide toxicity research [40–42]. Cell viability and proliferation was assessed by the mitochondrial-dependent reduction of MTT [3-(4,5-dimethylthiazol-2-yl)-2,5-diphenyltetrazolium bromide] to formazan using a MTT cell proliferation assay kit (Cayman Chemical, Item N0. 10009365). Confluent cells were incubated for 24 h incubation with the specified treatment. Caco-2 were exposed for 24 h to either the ternary mixtures or a two-chemical combination (i.e., combination of imidacloprid and nitrate, imidacloprid and atrazine or nitrate and atrazine). The next day 10 µL of MTT reagent was added to each well and mixed gently on an orbital shaker (Scientific Industries Mini-300 Orbital-Genie) for 1 minute. Following a 4 h incubation, 100 µL of crystal dissolving solution was added to each well and the plates were incubated for 4 h. Absorbance was read at 570 nm using a BioTek ELx808 microplate reader (BioTek Instruments Inc., Winooski, Vermont, USA). The results were expressed as % viability, which is the average absorbance of the wells/average absorbance of the control[43].

Three independent experiments were performed for ternary exposure experiments, and each treatment had 4 replicates. Four independent experiments with one replicate per condition were performed for the two-chemical exposure experiments. GraphPad Prism (Version 10.3.1) was used to perform a two-way ANOVA with Tukey’s post hoc test (P-value ≤ 0.05).

### 2.7 Live/Dead Caco-2 viability assay

The viability of Caco-2 cells was assessed with a live/dead assay. Confluent cells were seeded in 96-well plates and incubated for 24 h incubation with the specified treatment. A working solution of 200X Hoechst 33342 (Thermo Fisher, Lot: 2647883) and 2X propidium iodide (Invitrogen, Lot:359695-000) was prepared the day of the experiments. Aliquots of 11 µL of the working solution was added to each well and immediately imaged using a fluorescent microscope (Nikon Eclipse Ti) (20x magnification). Three independent experiments were performed, and each treatment had 3-5 replicates.

## 3. RESULTS

### 3.1 Ternary mixtures impact growth kinetics of poultry cecal microbiota

In the human gut microbiome, ingested xenobiotics interact with microbial enzymes within the small intestine and colon such as β-glucuronidases, nitroreductases, and sulfoxide reductases [44]. Analogous to human gut microbial composition, poultry gut microbiomes are predominated by *Firmicutes* and *Bacteroides* [44,45]. Given this similarity in gut microbial composition and that ingested contaminated groundwater, a xenobiotic, would be mediated within the gut microbiome [46], we selected a poultry cecal microbiome model that would provide insights into xenobiotic-mediated changes or other interactions within the cecal microbiome. Poultry cecal microbiome samples were exposed for 24 h to ternary mixtures as well as the environmentally relevant concentration of each chemical individually and growth kinetics for each treatment was observed to assess if cecal growth rate was lowered following exposure to mixtures of agricultural chemical. Overall, all treatments did not have a substantial lag phase (Figure 2). However, all treatment groups reached a similar maximum Optical Density over the 24 h time course (Figure 2). Growth rates of each treatment were then calculated to determine if there were differences in the exponential phase. It was observed that environmental concentrations of Imidacloprid (0.58 ppb), Nitrate (35, 000 ppb) and Atrazine (1.7 ppb) resulted in a growth rate of 0.1141 h^-1^, 0.1048 h ^-1^, and 0.1089 h ^-1^, respectively (Table 2). These are slower rates than ceca only growth rate of 0.1607 h ^-1^ but exceeds that of each ternary mixture, which had the slowest growth rates (Table 2). A similar trend was also observed for the equivalent and equipotent mixtures (Table 2). The presence of the mixtures decreased the growth rate by as much as 75%, while individual chemicals caused around a 30% decrease. The results demonstrate that ternary mixtures, even at environmentally relevant concentrations, substantially lower the growth rate of poultry cecal microorganisms.

**Figure 2.**
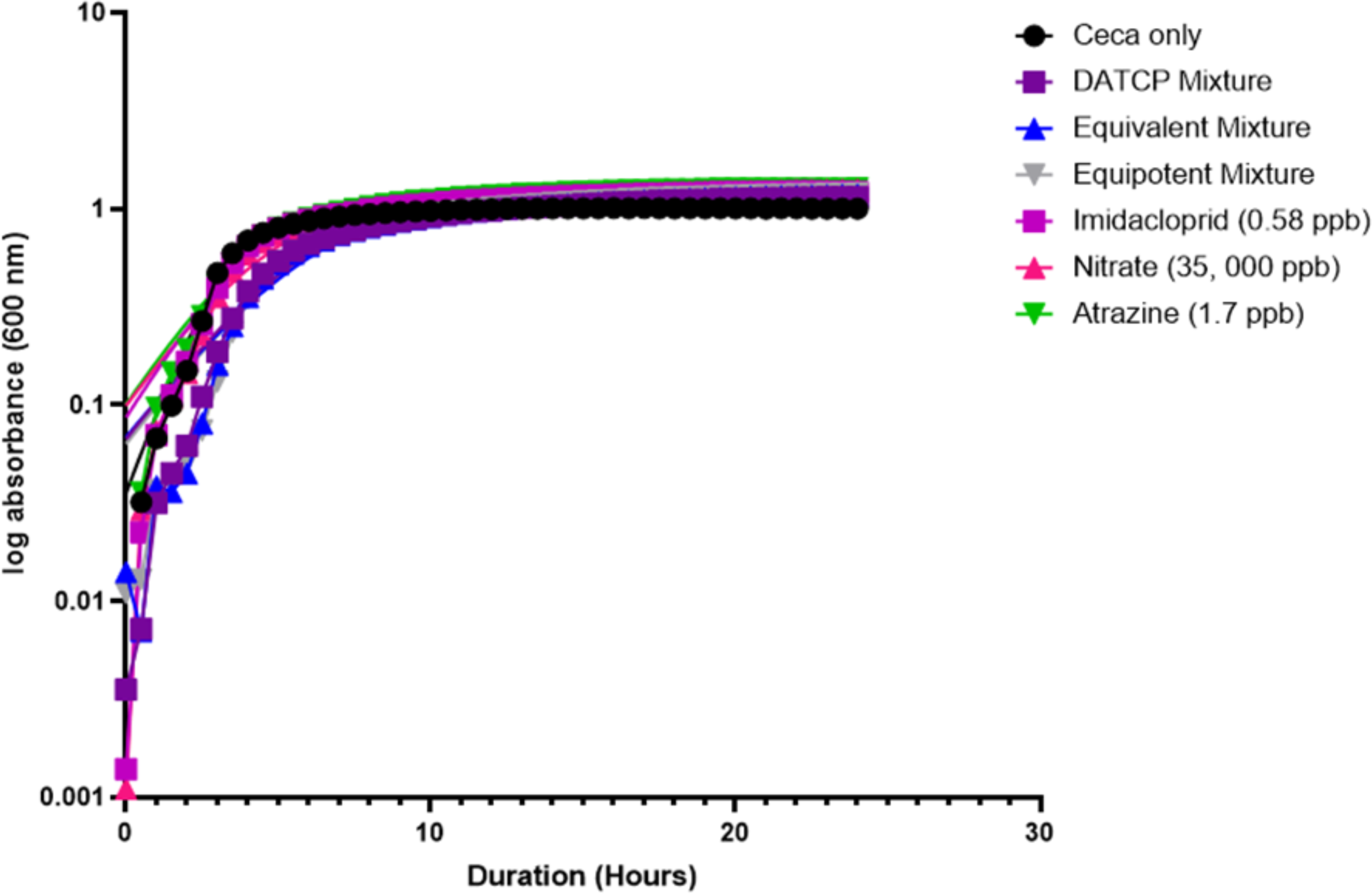
Growth curve of cecal microbiota following 24 h exposure to ternary mixtures and individual chemicals.

**Table 2.** Maximum growth rate of cecal microbiome cultures (h^-1^) during 24 h exposure to agricultural chemicals.

| <u>Treatment</u> | <u>Growth Rate</u> |
| --- | --- |
| Ceca only | $0.16 \text{ h}^{-1}$ |
| DATCP Mixture | $0.06 \text{ h}^{-1}$ |
| Equivalent Mixture | $0.06 \text{ h}^{-1}$ |
| Equipotent Mixture | $0.04 \text{ h}^{-1}$ |
| Imidacloprid (0.58 ppb) | 0.11 h<sup>-1</sup> |
| Nitrate (35, 000 ppb) | 0.10 h<sup>-1</sup> |
| Atrazine (1.7 ppb) | 0.11 h<sup>-1</sup> |

### 3.2 Caco-2 viability significantly impacted by two-chemical combinations

To assess the effects of the selected ternary mixtures on Caco-2 cell viability, a checkerboard method was incorporated. Caco-2 cells were exposed to a combination of imidacloprid and nitrate, imidacloprid and atrazine or nitrate and atrazine ranging from 3,000 ppb to 0 ppb (10-fold serial dilution). We predicted that higher concentrations of a two-chemical combinations (3,000 ppb of each chemical) would inhibit viability more than lower two-chemical combinations (30 ppb of each chemical). We did observe a significant decrease in Caco-2 cell viability for the two-chemical combinations (Figure 3; Tables S1-3). However, a two-chemical combination was not always required to elicit a significant impact on Caco-2 viability. For instance, 3,000 ppb nitrate singularly resulted in significantly lower cell viability compared to individual exposures at 30 ppb nitrate and 300 ppb nitrate (P-value ≤ 0.05; Figure 3A). Overall, it was observed that a two-chemical combination at the highest concentration (3,000 ppb) did yield a significant decrease in Caco-2 cell viability. For example, 3,000 ppb imidacloprid + 3,000 ppb atrazine results in significantly lower cell viability compared to the control, 300 ppb imidacloprid alone and 30 ppb imidacloprid alone (P-value ≤ 0.05; Figure 3B). Also, 3,000 ppb imidacloprid + 3,000 ppb atrazine resulted in significantly lower cell viability compared to 30 ppb imidacloprid + 300 ppb atrazine (P-value ≤ 0.05; Figure 3B). A similar trend was seen with 3,000 ppb nitrate + 3,000 ppb atrazine which significantly lowered cell viability compared to individual exposures to 30 ppb nitrate and 300 ppb nitrate (P-value ≤ 0.05; Figure 3C). In all, this data demonstrates the presence of a dose-response relationship.

**Figure 3.**
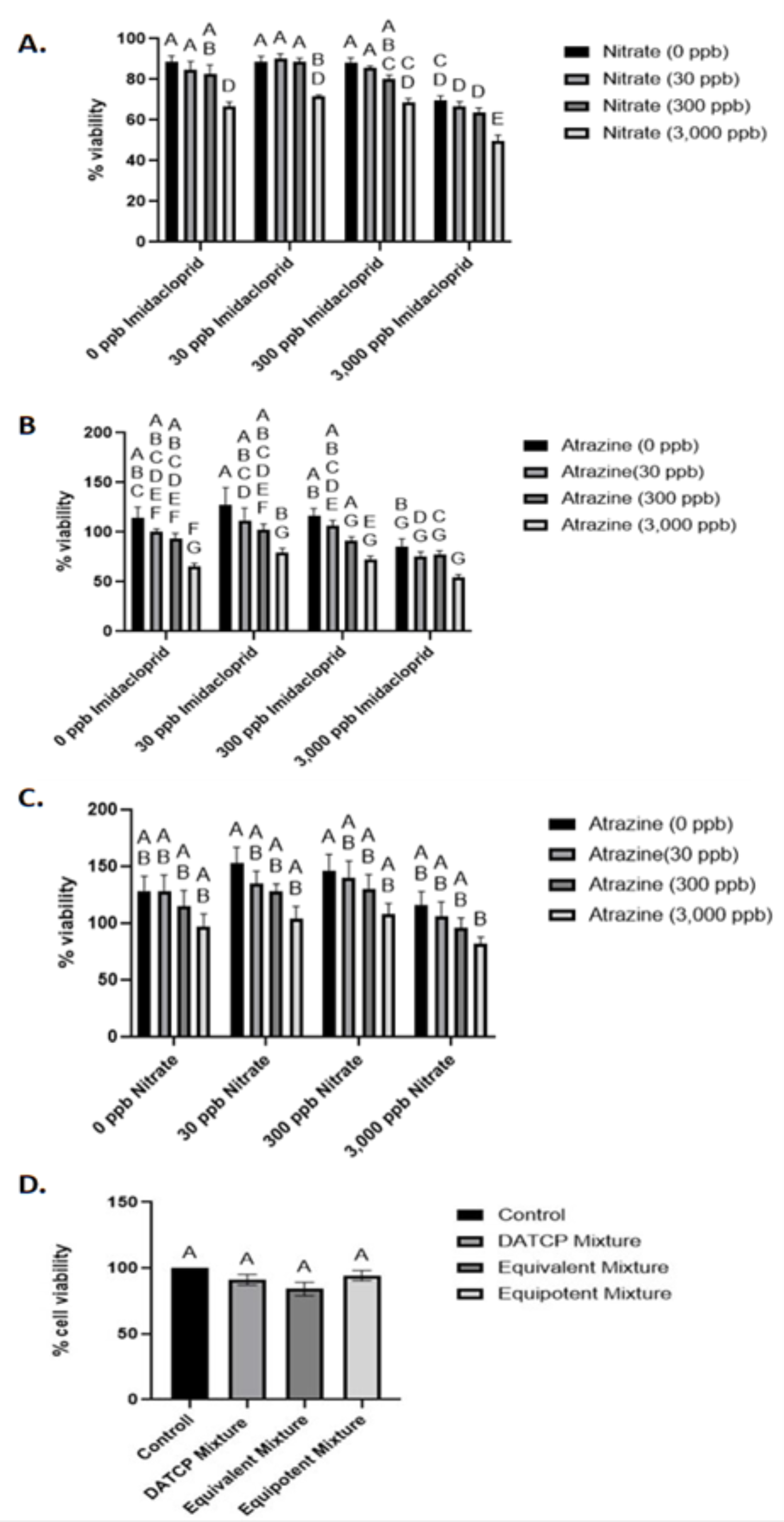
Caco-2 cell viability assays following exposure to two-chemical combinations and ternary mixtures of agricultural chemicals: (**A**) combinations of imidacloprid and nitrate, **(B**) combinations of imidacloprid and atrazine, and (**C**) combinations of nitrate and atrazine. Caco-2 cell viability was not significantly impacted following 24h exposure to ternary mixtures (**D**). Significance was determined with a two-way ANOVA (Tukey’s post-hoc test; P-value ≤ 0.05). A summary of the compact letter display is provided in supplemental tables 1-3.

Caco-2 cells were also exposed to three ternary mixtures over 24 to determine if environmental concentrations of nitrate, atrazine and imidacloprid decreased cell viability. Following a 24 h exposure, the results indicated that the DATCP mixture and equivalent mixture did result in lower viability compared to the control but were not statistically significant (Figure 3D). However, an acute exposure to the equivalent mixture decreased Caco-2 viability by approximately 16% (Figure 3D). Further, from confocal microscopy, we did not observe notable morphological changes to the Caco-2 cells that would indicate decreased viability following exposure to any ternary mixtures (Figures S3-S4). Thus, indicating that a 24 h exposure to the ternary mixtures did not significantly impact Caco-2 viability.

Given that the environmental concentrations did not result in complete death of the Caco-2 cells population, the minimum inhibitory concentrations were estimated (Figure S5). The measured viability of each chemical was plotted for each concentration tested and then fit with a linear regression. The predicted MIC, the x-intercept, was calculated, which estimates chemical concentration needed to elicit a cytotoxic effect. Based on this prediction it was determined that the environmentally relevant concentrations used as mixtures in our tested conditions are orders of magnitude below concentrations needed to completely inhibit Caco-2 viability.

### 3.3 *In silico* methods prioritized chemicals and chemical mixtures of concern

To determine if the water quality data used in this study corresponds with known chemical-biological interactions, an *in silico* tool, *toxEval,* was used to calculate exposure-activity ratios for each detected chemical. EARs above the threshold of 10^-3^ correspond to potential biological effects, as described in the methods. An EAR > 1 indicates the chemical concentration in the data set exceeds the chemical concentration determined to elicit a biological effect in a ToxCast high-throughput assay, which are derived from the US EPA’s chemical characterization database, ToxCast. An examination of the maximum EAR by chemical indicates that groundwater wells tested in Waushara and Richland were the only counties with max EARs between 0.06 and 0.22, which exceed the EAR threshold for concern (EAR threshold = 10^-3^) (Figure 4).

**Figure 4.**
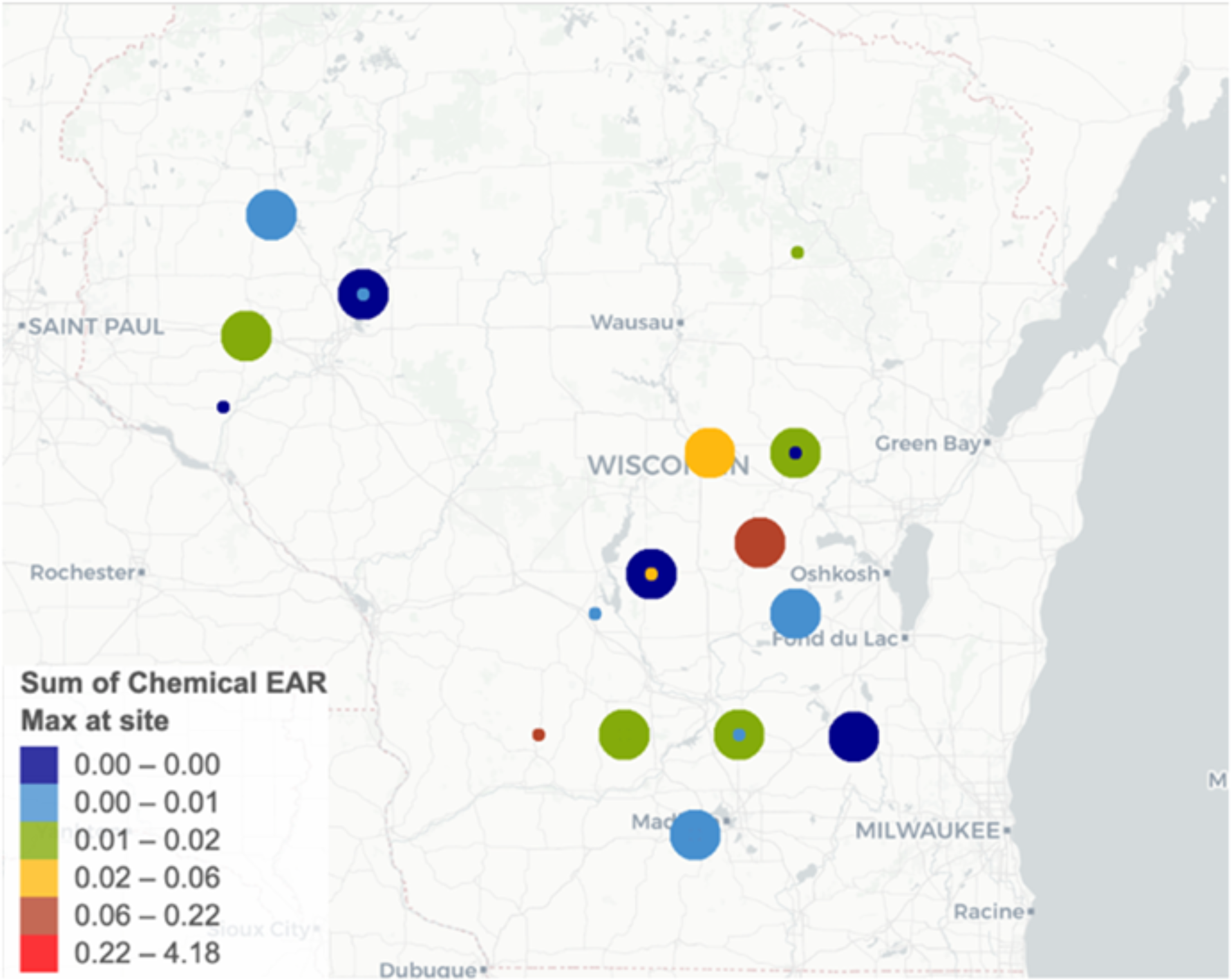
Summary of maximum exposure activity ratios by chemical within each sampling location (N = 16).

The maximum EAR for each prioritized chemical (EAR ≥ 10^-3^) was subsequently evaluated by chemical class to assess if the presence of a given class of pesticides (i.e., herbicides, insecticides and fungicides) was more strongly correlated with potential biological effects. We observed that the herbicides imazapyr, bentazone, alachlor, metachlor, atrazine, clomazone, fomesafen, and flumetsulam had reported EAR ≥ 10^-3^ (Figure 5). In addition, one fungicide, metalaxyl, was prioritized (EAR threshold = 10^-3^) (Figure 5). There were several ToxCast endpoints, representing negative biological effects from various assay results in a public database, corresponding to these prioritized chemicals (EAR > 10^-3^) (Figure 6). The chemicals included: bentazone, atrazine, atrazine TCR, fomesafen, metachlor, and alachlor (Table S7; Figure 6).

**Figure 5.**
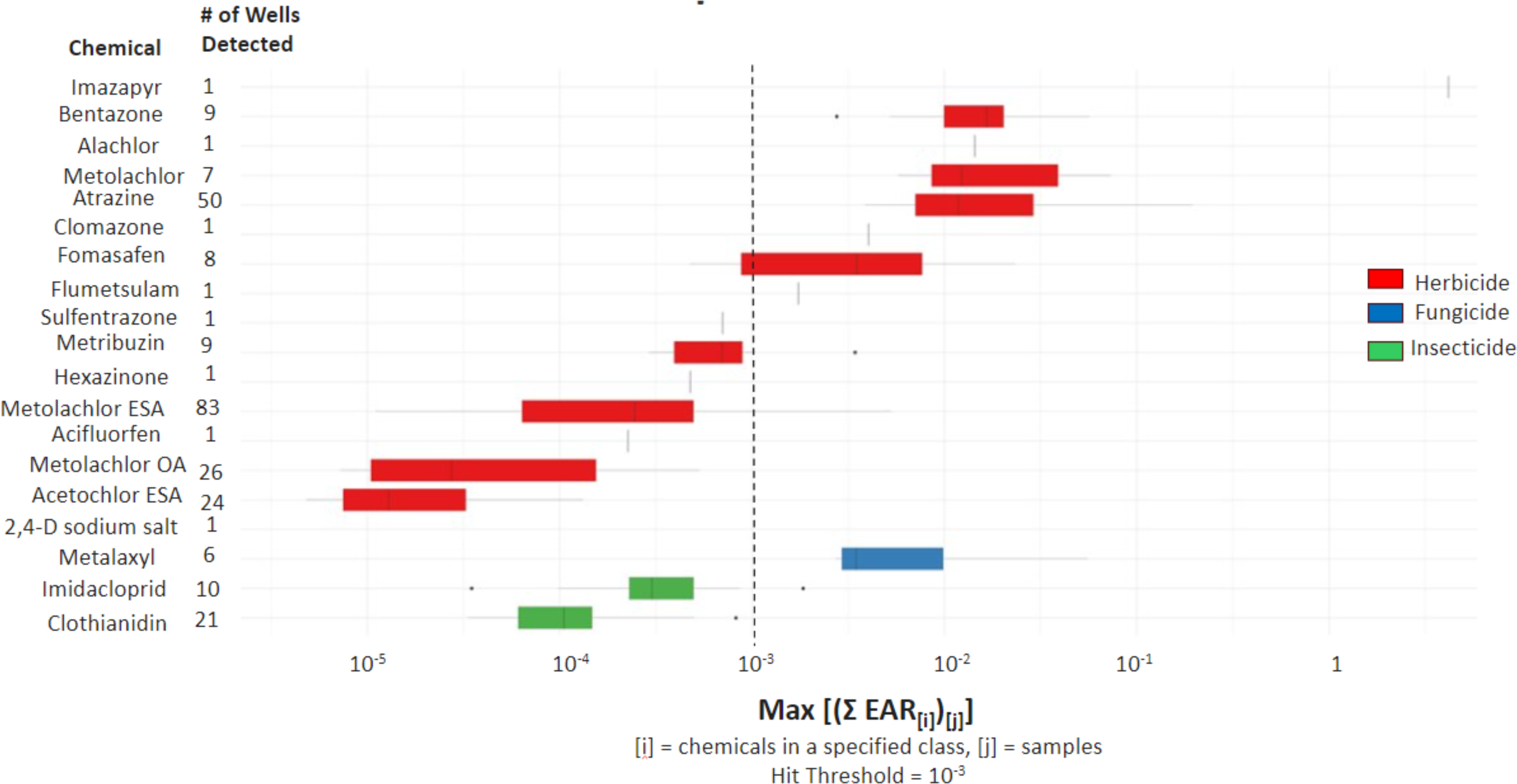
Summary of maximum exposure activity ratios for herbicides (red), fungicides (blue), and insecticides (green). A total of nine chemicals detected in the 2021 DATCP Targeted Sampling Report were prioritized (EAR ≥ 10^-3^).

**Figure 6.**
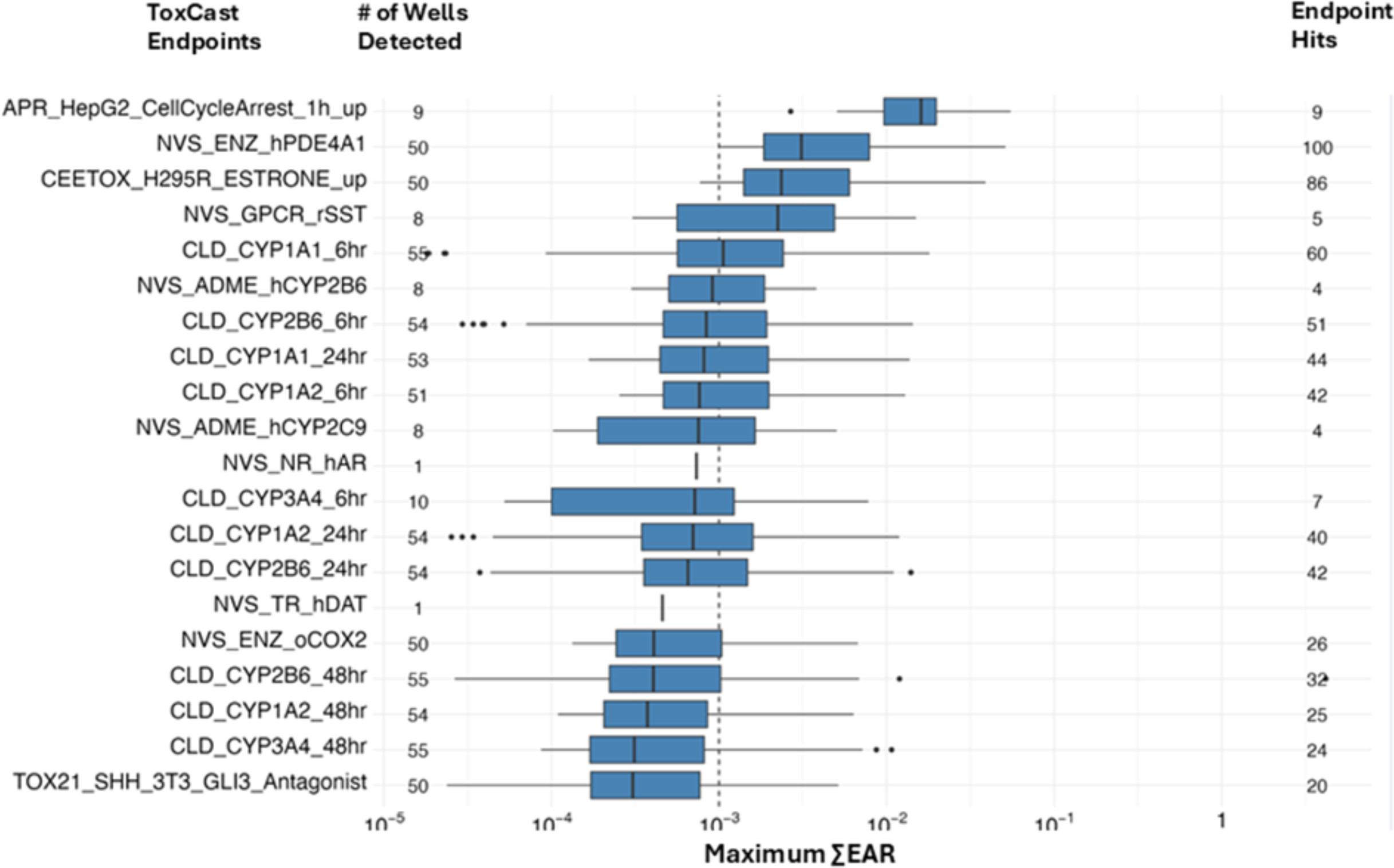
Summary of maximum exposure activity ratios for chemicals and corresponding ToxCast assays. ToxCast assays with EAR ≥ 10^-3^ indicate potential risk for biological risks for the chemical(s) detected in the endpoint hits.

Bentazone corresponded to the endpoint, APR_HepG2_CellCycleArrest_1h_up, a test to measure perturbation to cell phenotypes (i.e., changes to morphology or cell signaling) (Table S7; Figure 6). Atrazine and atrazine TCR corresponded to NVS_ENZ_hPDE4A1 (gene activity assay), CEETOX_H295R_ESTRONE_up (hormone induction assay) and CLD_CYP1A1_6hr (mRNA induction assay), which are tests of hormone synthesis, enzymatic activity and xenobiotic metabolism (Table S7; Figure 6). Fomesafen was associated with NVS_GPCR_rSST (radioligand binding assay) (Table S7; Figure 6). A summary of all detected endpoints by site is provided in Table S8. This data suggests that these chemicals have bioactivity at levels observed in some Wisconsin groundwater wells given that similar concentrations resulted in phenotypes correlated to adverse outcome pathways, as predicted with *toxEval* and ToxMixtures.

The *in silico* tool, ToxMixtures, was employed to evaluate the impact of chemical mixtures within this water quality data for potential chemical-biological effects. As done with the individual chemical prioritization, chemical concentration data from the 2021 Wisconsin DATCAP groundwater report was screened. The results indicated that two sampling locations, groundwater well DB719 (Waupaca) and groundwater well EW833 (Chippewa) contained chemical mixture concentrations which correlated to ToxCast assay endpoints, genes, and adverse outcome pathways (Table 3). However, groundwater well EW833 only contains atrazine which corresponded to gene target, PDE4A inhibition (Table 3). Groundwater well DB719 contained metolachlor ESA, metalaxyl, metolachlor OA, imidacloprid and atrazine (Table 3). The chemical mixture in groundwater well DB719 corresponded to several gene targets including CHRNA2 inhibition, CYP2B6 activation and CYP3A4 activation (Table 3). Adverse outcome pathways associated with the chemical mixture in groundwater well DB719 include upregulation of thyroid hormone catabolism, percellome toxicogenomic approach for AOP building, NR1I2 activation leading to hepatic steatosis, AhR activation leading to early life stage mortality, and cyclooxygenase inhibition leading to reproductive failure or dysfunction (Table 3). Overall, the prioritized chemical mixture was associated with 9 gene targets and 11 adverse outcome pathways (Table 3). These results, both from *in vitro* and *in vivo*, suggest that environmentally relevant exposure levels may have associated negative health effects.

**Table 3.** Summary of sampling locations with chemical mixtures linked to adverse outcome pathways as determined with ToxMixtures.

| Sampling Location | Chemical | Gene symbol | Summary of Adverse Outcome Pathways (AOPs) |
| --- | --- | --- | --- |
| DB719 - Waupaca | Metolachlor ESA | CHRNA2_Inhibition | Upregulation of Thyroid Hormone Catabolism |
|  | Metalaxyl | CYP2B6_Activation | Percellome Toxicogenomic Approach for AOP Building |
|  |  | CYP3A4_Activation |  |
|  | Metolachlor OA | NR1I2_Activation | NR1I2 activation leading to hepatic steatosis |
|  | Imidacloprid | CYP1A1_Activation | AhR activation leading to early life stage mortality |
|  |  | CYP1A2_Activation |  |
|  | Atrazine | Esterone_Activation | Cyclooxygenase inhibition leading to reproductive failure |
|  |  | PDE4A_Inhibition | Cyclooxygenase inhibition leading to reproductive dysfunction |
|  |  | PTGS2_Inhibition |  |
|  |  |  | Gastric ulcer formation |
| EW833 - Chippewa | Atrazine | PDE4A_Inhibition | Not Available |

## 4. DISCUSSION

In this study, we aimed to elucidate chemical-biological interactions and adverse outcome pathways associated with an environmentally relevant mixture of agricultural chemicals detected in Wisconsin groundwater. We selected nitrate, atrazine and imidacloprid based on their frequency of detection and concentrations exceeding enforcement standards in the 2021 DATCP Targeted Sampling Report. There are several challenges with evaluating mixtures. Some of which include the abundance of data for a limited number of chemicals and the inability to accurately account for all realistic exposures in a single experiment [30]. Therefore, the most appropriate method is to assess ecologically relevant chemical mixtures at various concentrations. This method combination provided a detailed account of acute exposure effects on the primary exposure route in a body but also a prediction of the effects of chronic exposure on other systems. The incorporation of *in vitro* and *in silico* methods in this study also provided a reproducible and effective method that could be utilized yearly to evaluate future groundwater quality data across the state.

In this study, we first aimed to elucidate the effects of two-chemical combinations and ternary mixtures of nitrate, atrazine and imidacloprid on two models, poultry cecal microorganisms and Caco-2 cells. There were clear differences in eukaryotic versus prokaryotic interactions with these agricultural chemicals. It is well documented that the host gut microbiota interacts with xenobiotics via direct and indirect mechanism(s) which can impact host health [44,46]. Typically, a xenobiotic such as an environmental toxicant would interact with the intestine and colon microbiota, if they are not absorbed into the gastrointestinal tract and transformed upon ingestion following interaction with microbial enzymes [44]. Xenobiotics may also cycle back into the gut through enterohepatic circulation. In this process the xenobiotic is first synthesized by the liver and subsequently metabolized by the gut microbiota [44,47].

To our knowledge there are no current studies that have explored if the metabolism of a mixture of nitrate, atrazine and imidacloprid alters each chemical’s normal metabolism in the gut microbial ecosystem. Current literature has demonstrated that oral microorganisms [48–51] and gut microorganisms [52–55] are capable of nitrate metabolism which in turn, affects the health of the host. Imidacloprid is commonly degraded via oxidation [56]. However, there are several reported imidacloprid-degrading microorganisms [57–59]. Fungal and bacterial communities have also been shown to mineralize atrazine [60–62]. For instance, Henn et al.(2020) [60] discovered that fungal strains, *Pluteus cubensis* SXS320, *Gloelophyllum striatum* MCA7, and *Agaricales* MCA17 were capable of depleting 30-38% of total atrazine in a liquid culture. Interestingly, they noted that fungal degradation of atrazine relies on nitrogen availability [60]. Atrazine has also been demonstrated to degrade in the presence of bacterial communities [61,62]. Billet et al. (2019) [61] demonstrated the ability of an evolved bacterial community of *Chelatobacter* sp. SR3822, *Pseudomonas* sp. ADPe19, *Arthrobacter* sp. TES23 and *Variovorax* sp. 38R to degrade atrazine. However, degradation of atrazine was lower in the presence of organic nitrogen sources, ammonium and cyanuric acid [61]. Yang et al.(2010) [62] demonstrated that 83.3% of atrazine can be degraded by a community *Klebsiella* sp. A1 and *Comamonas* sp. A2 but noted lower levels of degradation in the presence of (NH_4_)_2_SO_4_ and NH_4_NO_3_. The chemical mixtures tested in this study will likely have varied degrees of degradation within an anaerobic environment like a gastrointestinal tract. Furthermore, given that most bacteria capable of degrading atrazine can use atrazine as a nitrogen source [62], it is plausible that inorganic and organic nitrogen sources will have varied impacts to the biodegradation of atrazine in the gut microbiome. These complex biotic and abiotic interactions could provide some explanation for differences in observed phenotypes from two-chemical and ternary exposure experiments in this study. Further, given that our tested mixtures did not significantly decrease cell viability, cytotoxic evaluation is not the most valuable metric to use for understanding the chemical-biological interactions that may occur following ingestion of agricultural chemicals at environmentally relevant concentrations.

In this study Caco-2 viability was lower when exposed to ternary mixtures but not significantly different from controls, but a more pronounced decrease in Caco-2 viability was observed following exposure to two-chemical combinations. However, an acute exposure to the equivalent mixture decreased Caco-2 viability by approximately 16% (Figure 3D). It is possible that interaction of the three chemicals result in an antagonistic effect. Additionally, only Caco-2 cells exposed to 50 µM atrazine and 25 µM atrazine impacted proliferation and viability [63]. These results highlight the resilience of this human epithelial cell line to agricultural toxicants. In addition, it is possible that the chemicals are being biotransformed following interaction with the host or independent of the host. Based on the observations in this study, it is plausible that the cumulative effects of small decreases in cell viability over the course of a chronic exposure to the environmentally relevant mixtures and two-chemical combinations would alter intestinal barrier function, which may subsequently impair nutrient uptake in the gastrointestinal tract or increase incidence of tumorigenesis.

In this study, *in silico* methods were incorporated for further evaluation of the water quality data. The inclusion of these tools provided predictions on chemical bioactivity. We noted the presence of five prioritized chemicals within two groundwater wells which corresponded to nine genes (Table 3). The first, CHRNA2, is a mammalian gene that encodes the nicotinic acetylcholine receptors (nAChRs) [36]. This gene target is likely correlated to the presence of imidacloprid in this groundwater well as this neonicotinoid acts on the nAChRs in insects [36,64]. Notably, this groundwater well is also linked to genes CYP2B6, CYP3A4, CYP1A1 and CYP1A2 which are encoded in the cytochrome P450 superfamily of enzymes. Cytochrome P450 enzymes are involved in xenobiotic metabolism [65]. Based on the observed data, atrazine is the only chemical detected in groundwater well DB719 which was linked to ToxCast assays associated with these gene targets (Table 3; Table S8). In addition, atrazine presence was associated with gene, PDE4A (Table 3). PDE4A is a signal transducer that is vital in regulating cAMP [66]. High expression of this isoform of PDE4 has also been documented in cancers [67] and has been considered a marker of carcinogenesis [67–69]. Other gene targets included the gene NR1I2, a transcription factor that aids in detoxification [36] and esterone activation.

These results demonstrate the possibility for a reproducible method for assessing implications of groundwater water quality data. The evaluation of an acute exposure to an ecologically relevant ternary chemical mixture of nitrate, atrazine and imidacloprid was demonstrated to exhibit varied biological impacts to poultry cecal microbiota and Caco-2 cells which demonstrates the potential sensitivity of the gut microbe to chemical toxicity. In addition, the altered growth rate of the microbiome samples and decreased Caco-2 viability indicates that chronic exposure may result in significant modulation to microbial activity and intestinal permeability. Further, the *in silico* tools provided a means of determining which chemicals are of highest potential concern based on similarities in chemical concentrations exhibited to cause chemical-biological interactions. Coupled with the results from the biological assays performed in this study, there is sufficient evidence that a mixture of nitrate, atrazine and imidacloprid, even at environmental concentrations, adversely impacts gut microbiota and gut epithelial cells.

## 5. CONCLUSIONS

In summary, we observed chemical-biological interactions for both models utilized, poultry cecal microbiota and Caco-2 cells, following an acute exposure to a mixture of nitrate, atrazine and imidacloprid. It was observed that exposure to the ternary mixtures resulted in a decline in growth rate for poultry cecal microbiome samples which could contribute to dysbiosis in the poultry cecal microbiome. The same ternary mixtures did not significantly decrease Caco-2 viability, but two-chemical combinations of the agricultural chemicals did significantly decrease Caco-2 viability demonstrating either antagonistic effects or chemical biotransformation following interaction with the host or among the chemicals independent of the host. These results emphasize the importance of elucidating the impacts of chemical mixtures on various models and further imploring *in silico* methods to predict potential adverse effects for communities utilizing this groundwater. Doing so will allow for continuous and reliable evaluation of water quality data collected from Wisconsin groundwater wells.

## DATA AVAILABILITY

Data are available within the manuscript. Additional data are provided in the supplemental sections. Raw data are available upon request.

## Supporting information

Supplemental Figures_GWIV

## ACKNOWLEDGEMENTS

The authors would like to thank Dr. Ophelia Venturelli and Julie Duclos for providing the Caco-2 cells used in this study and offering cell culture training. We would also like to thank Dr. Steve Ricke for providing access to various instruments used in this study. The author CC would like to thank the University of Wisconsin-Madison’s Molecular and Environmental Toxicology graduate program for its financial support through the Molecular and Environmental Toxicology Training Award (National Institute of Environmental Health Sciences; T32 ES007015) and the Science and Medicine Graduate Research Fellowship for their salary financial support.

## AUTHOR CONTRIBUTION STATEMENT

The author ELW was responsible for study conceptualization, methodology development, project administration and supervision and writing – reviewing and editing. The author CC was responsible for formal analysis, investigation, data visualization and writing – original draft preparation.

## CONFLICT OF INTEREST

No, there is no conflict of interest.

## SUPPLEMENTAL MATERIALS

Supplemental Tables S1-8

Supplemental Figures S1-5

## Notes

### Competing Interest Statement

The authors have declared no competing interest.

