## Supplemental Figures_GWIV for "Acute exposure to groundwater contaminants mixture of nitrate, atrazine and imidacloprid impacts growth kinetics of poultry cecal microbiomes and significantly decreases Caco-2 cell viability"

**Supplementary Tables 1-8 are in Excel fille**

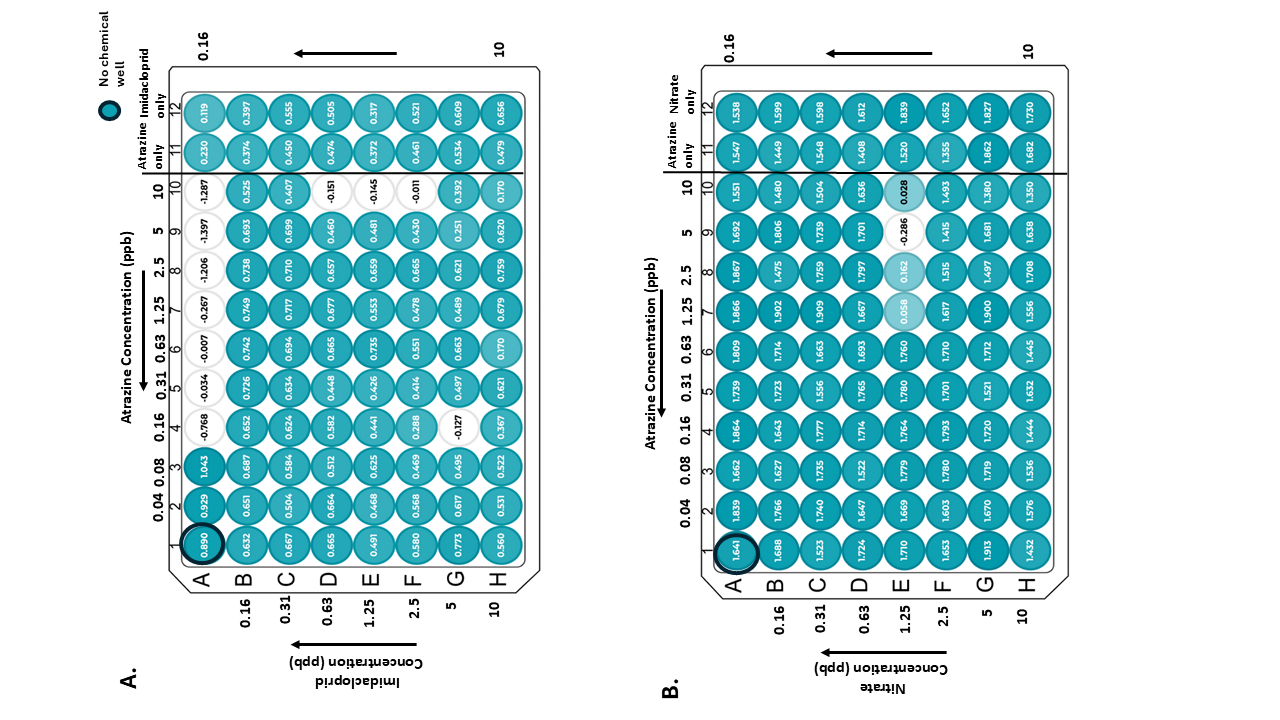

Figure S1. Absorbance (OD_600 nm)_ of poultry cecal samples. Growth kinetic experiment using a checkerboard method to assess A) a two-chemical combination of atrazine and imidacloprid (300 to 0.16 ppb) and B) a two-chemical combination of nitrate and atrazine (300 to 0.16 ppb)

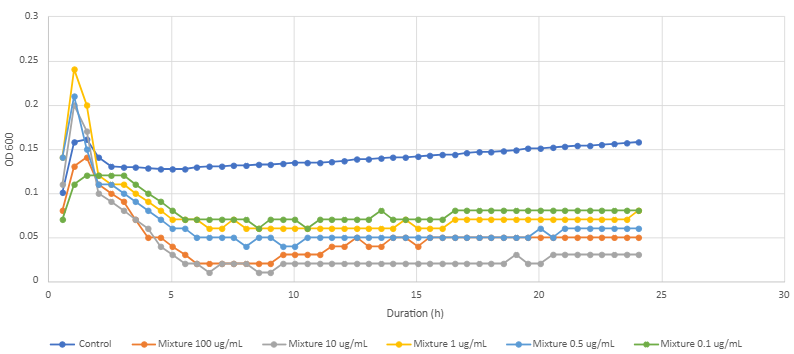

Figure S2. Growth curve of poultry cecal microbiome exposed to ternary mixtures at equivalent concentrations ranging from 100 µg/ml (100 ppb) to 0.1 (0.1 ppb). Poultry cecal microbiomes grown in Luria-broth at 37**°**C for 24 h. Absorbance read at OD_595 nm_.

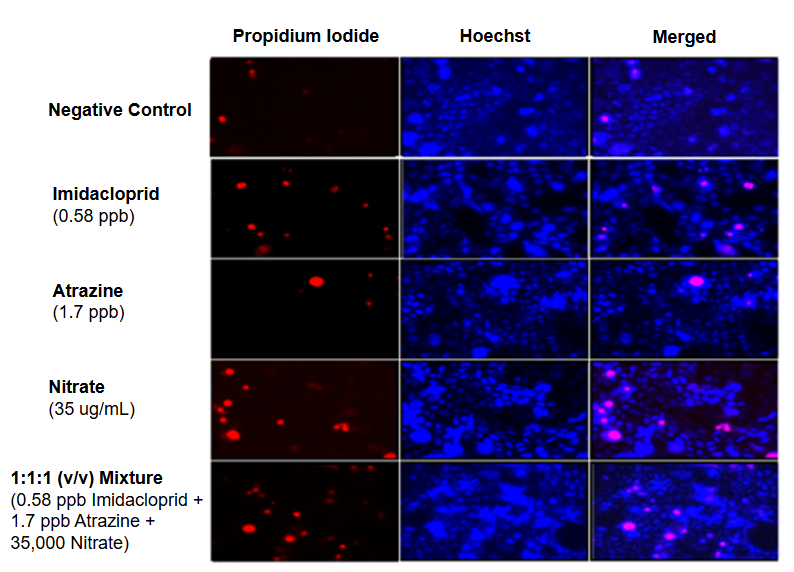

Figure S3. Live/Dead staining of Caco-2 cells exposed to environmentally relevant concentrations of nitrate, atrazine and imidacloprid as well as a ternary mixture of the chemicals for 24 h.(20x magnification).

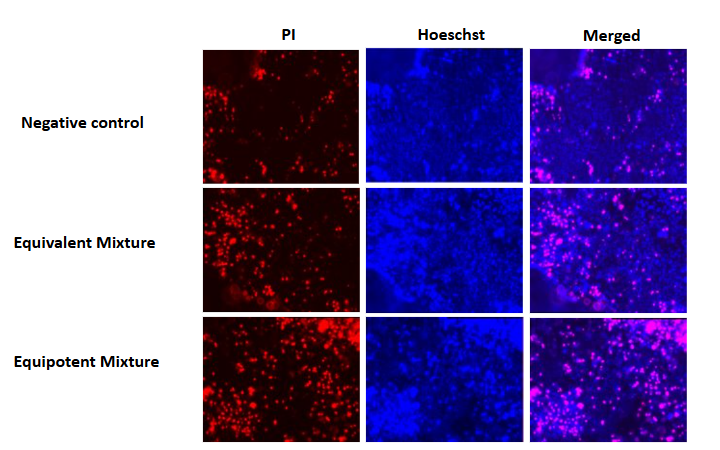

Figure S4. Live/Dead staining of Caco-2 cells exposed to the equivalent mixtures (100,000 ppb nitrate + 100,000 ppb atrazine + 100,000 ppb imidacloprid) and the equipotent mixtures (1,000 ppb nitrate + 10,000 ppb atrazine + 10,000 ppb imidacloprid) for 24 h (20x magnification).

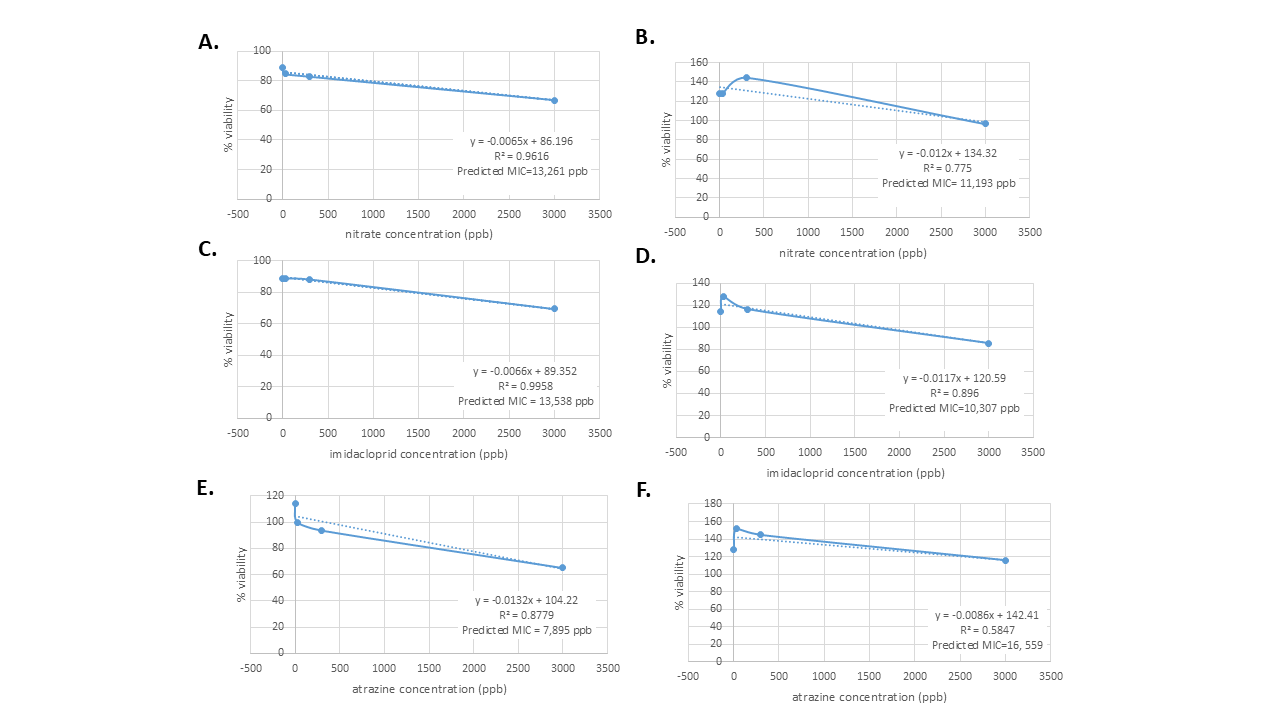

Figure S5. The percent viability for Caco-2 cells exposed to two-chemical combinations was used to predict the minimum inhibitory concentration (MIC) for each chemical. The slope of each chemical was first plotted, and the predicted MIC (“x”) was subsequently calculated. The results were as follows: A) the predicted MIC of nitrate when combined with imidacloprid, B) the predicted MIC of nitrate when combined with atrazine, C) the predicted MIC of imidacloprid when combined with nitrate, D) the predicted MIC of imidacloprid when combined with atrazine, E) the predicted MIC of atrazine when combined with imidacloprid and F) the predicted MIC of atrazine when combined with nitrate.
